## Supplemental files for "A bioengineered human urothelial organoid model reveals the urine-urothelium interplay in tissue resilience and UPEC recurrence in urinary tract infections"

### Supplementary Information

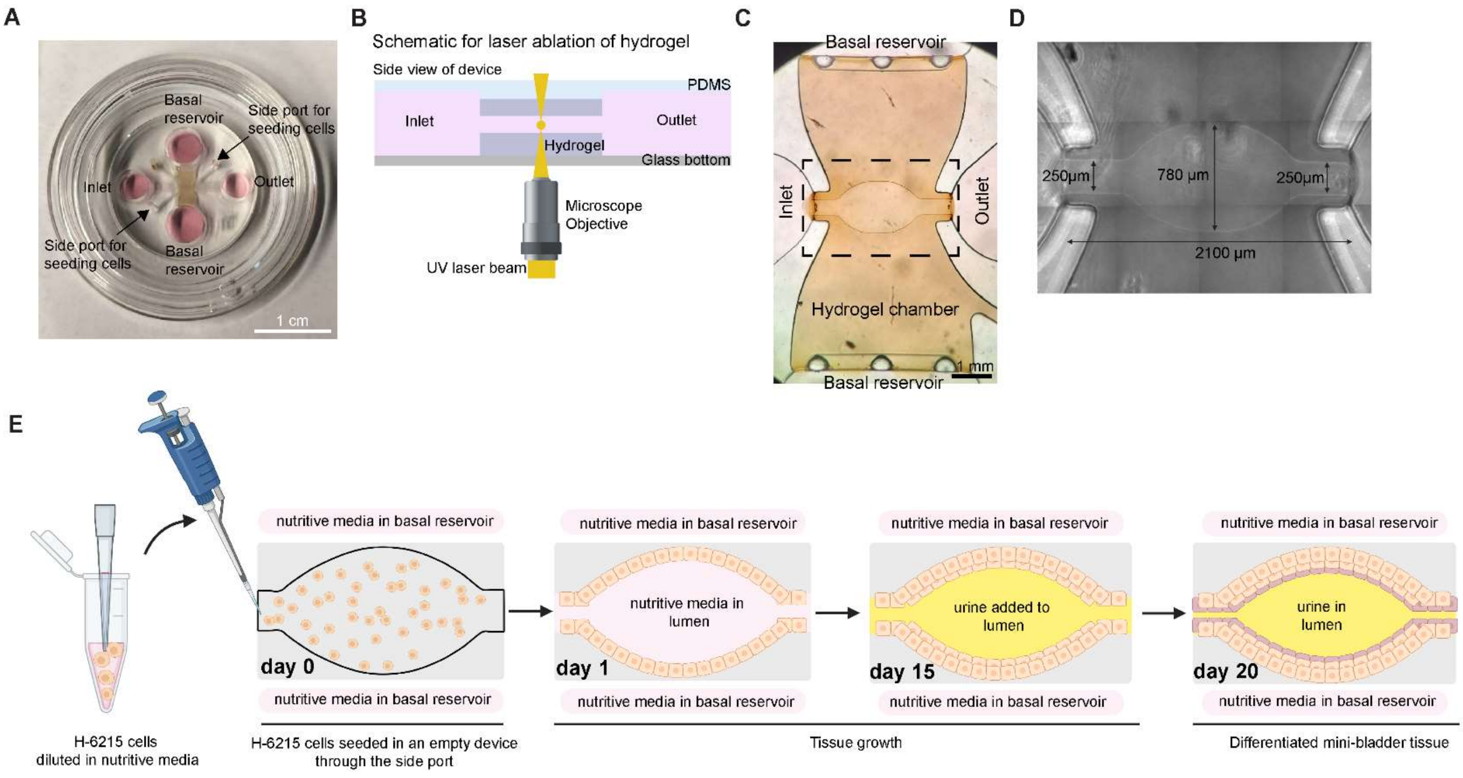


**Figure S1: Human mini-bladder architecture and tissue culture timeline**

(**A**) Photograph of a mini-bladder device, scale bar = 1 cm. **(B)** Schematic cross-sectional view of the device as positioned during ablation by the 2-photon laser. **(C)** Hydrogel chamber with the urine channel (highlighted by the black dashed box) in the central chamber after laser ablation **(D)** An optical image of the mini-bladder form generated after laser ablation, the dimensions of the urine channel and bladder are overlaid. **(E)** Schematic representation of the timeline of tissue culture.

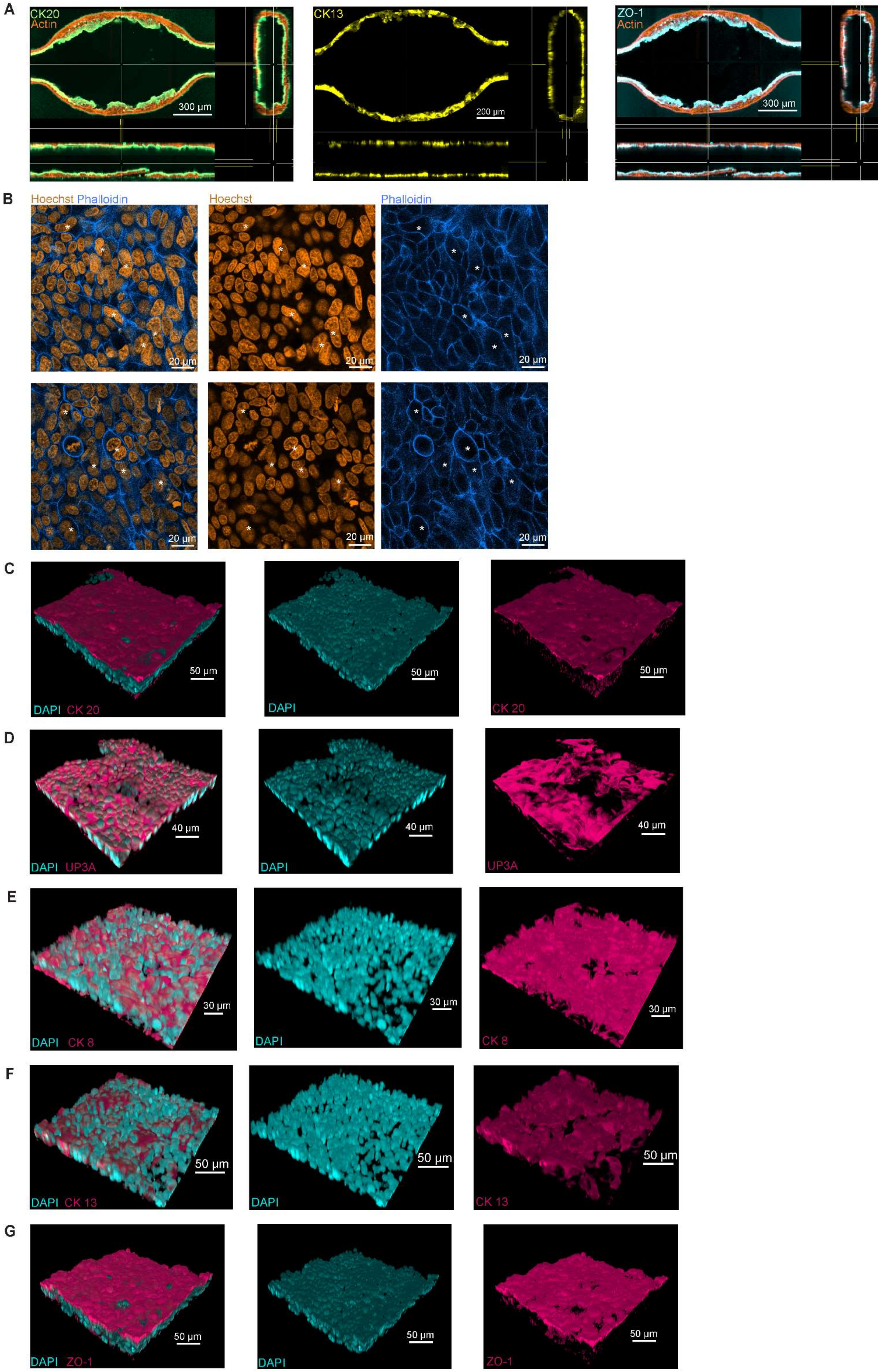
A
CK20
Actin
300 µm
CK13
200 µm
ZO-1
Actin
300 µm
B
Hoechst Phalloidin
Hoechst
Phalloidin
20 µm
C
DAPI CK 20
DAPI
CK 20
50 µm
D
DAPI UP3A
DAPI
UP3A
40 µm
E
DAPI CK 8
DAPI
CK 8
30 µm
F
DAPI CK 13
DAPI
CK 13
50 µm
G
DAPI ZO-1
DAPI
ZO-1
50 µm

**Figure S2: High magnification characterization of urothelial architecture in mini-bladder**

(**A**) Intensity adjusted images corresponding to **Fig 1 C, E, G** showing orthogonal view of microtissue with CK20 (green), CK13 (yellow) and ZO-1 (cyan) respectively. **(B)** Representative images show binucleated, honeycomb- like cells in white asterisks (actin, azure; Hoechst, amber). (**C -F**) 3D views of confocal images of magnified sections of mini bladder shows CK20 (**C**), UP3A (**D**), CK8 (**E**), CK13 (**F**), ZO-1 (**G**) in magenta; Hoechst in cyan.

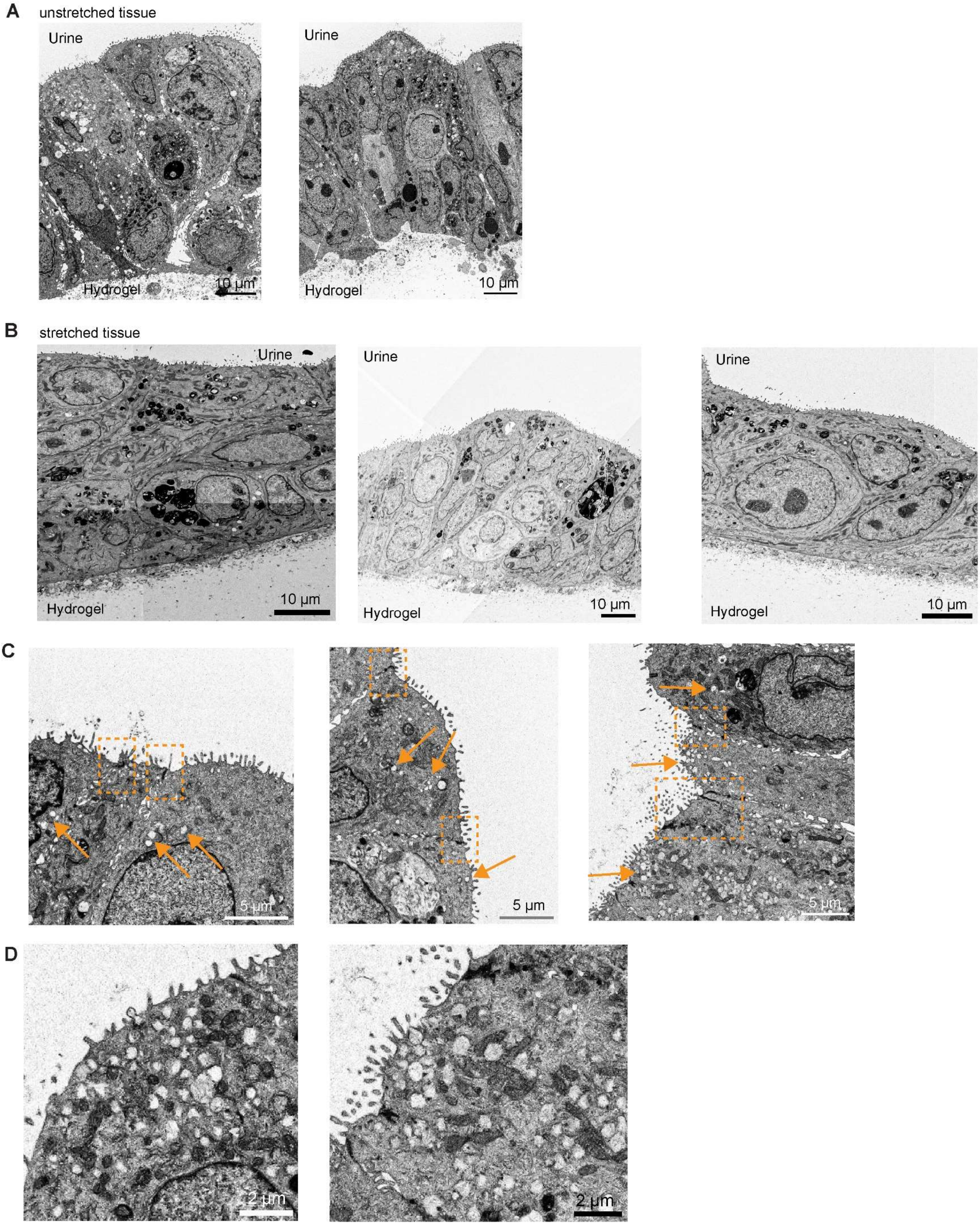
A
unstretched tissue
Urine
Hydrogel
10 µm
Urine
Hydrogel
10 µm
B
stretched tissue
Urine
Hydrogel
10 µm
Urine
Hydrogel
10 µm
Urine
Hydrogel
10 µm
C
5 µm
5 µm
5 µm
D
2 µm
2 µm

**Figure S3: Array tomography SEM images of unstretched and stretched tissue cross-sections**

**(A-B)** Images from array tomography SEM of unstretched (**A**) and stretched (**B**) tissue cross-sections**.** **(C)** Magnified images show vesicles (amber arrows) and tight junctions (amber dashed boxes) between lumenal cells in unstretched tissue. **(D)** Zooms show several vesicles in lumenal cells in unstretched tissue.

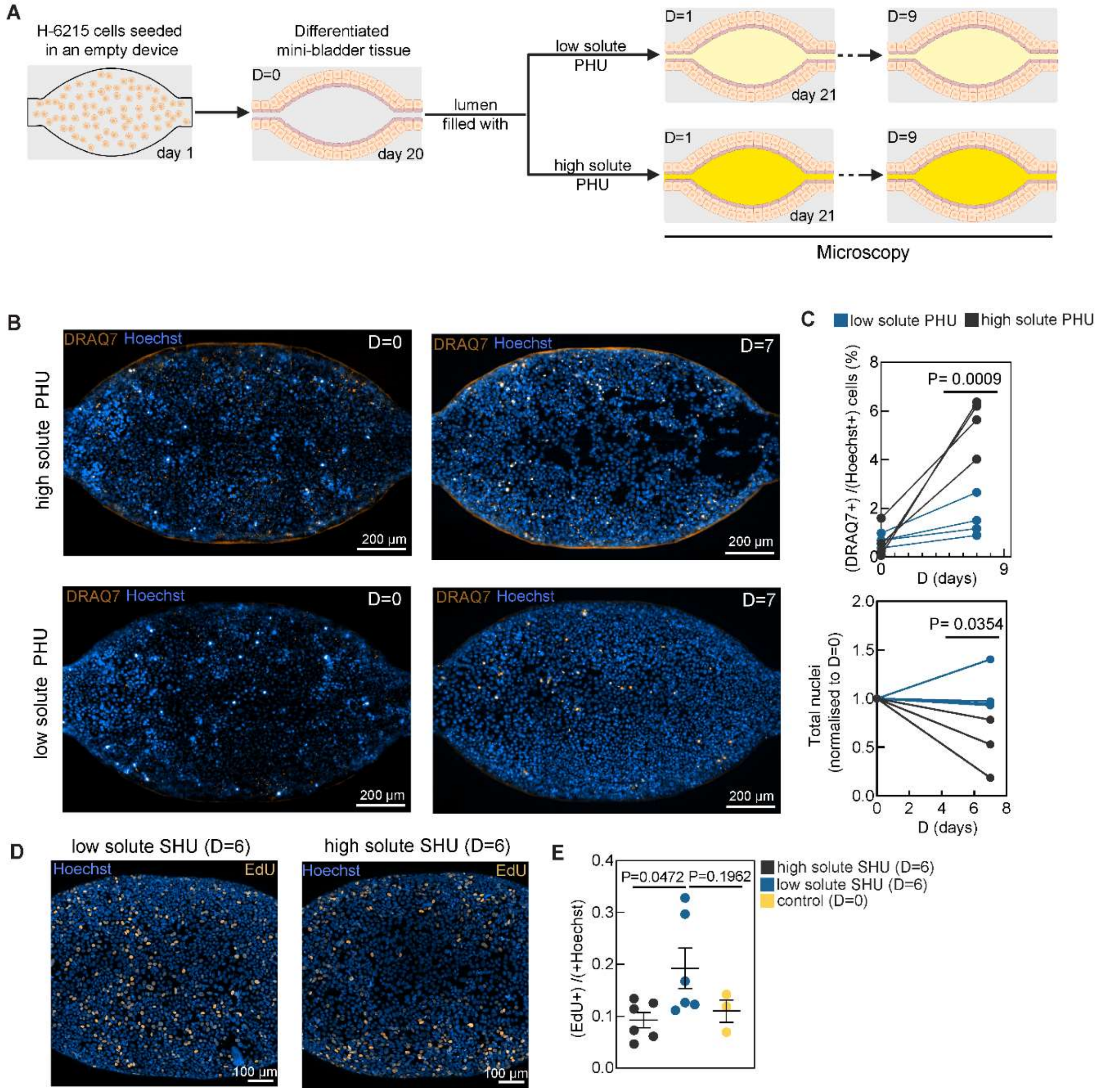


**Figure S4: Long term exposure to high solute concentration pooled human urine causes tissue damage**

**(A)** Timeline of long-term exposure of differentiated mini-bladders (D=0) to low and high solute concentration PHU for D=9 days. **(B)** Snapshots (maximum intensity projection) from timelapse imaging of mini-bladders in high and low solute concentration PHU (Hoechst in blue, DRAQ7 in amber). **(C)** Frequency of dead cells (DRAQ7+ nuclei) and total cell numbers (Hoechst+) over time for mini-bladders in high (n=4; 3 respectively) and low (n=4; 4 respectively) solute SHU. **(D)** Representative images of mini-bladders exposed to high and low solute SHU for D=6 (EdU in amber, Hoechst in blue). **(E)** Quantification of EdU+ nuclei in tissue exposed to high (n=6) and low (n=6) solute SHU for D= 6 and control tissue at D=0 (n=3). Data represented as mean ± SEM. P-values calculated using unpaired t test in (**C**), one-way Anova with Dunnett's multiple comparison test in (**E**).

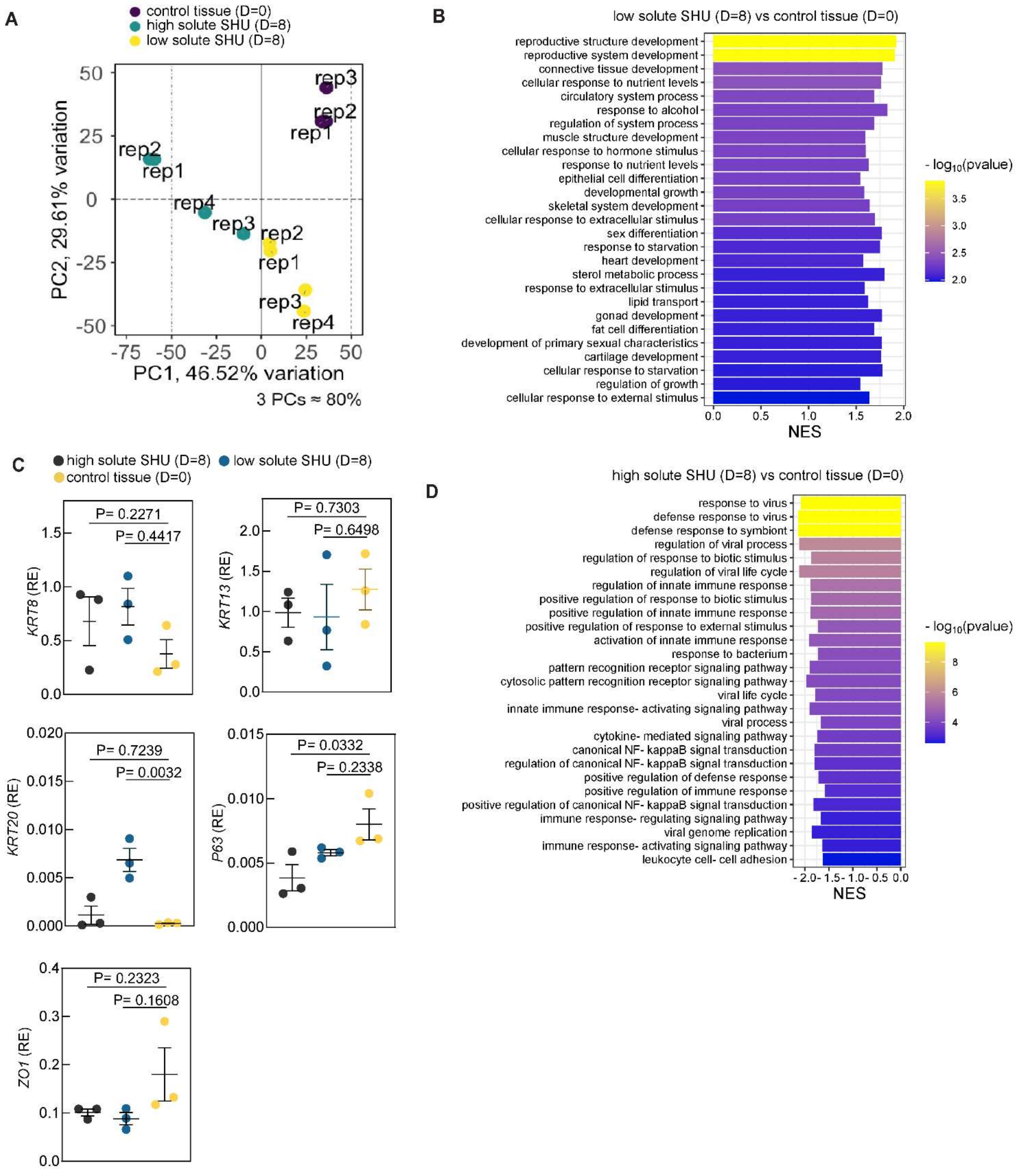


**Figure S5: Transcriptional profile of mini-bladder exposed to high and low solute SHU at D=8 versus control tissue at D=0**

**(A)** Principal component analysis (PCA) plot of mini-bladders exposed to high solute SHU for D=8 (n=4), mini-bladders exposed to low solute SHU for D=8 (n=4) and mini-bladders at D=0 (n =3) namely control tissue. **(B)** Top positively enriched GO BP pathways ordered by significance in low solute SHU exposed tissue at D=8 vs. control tissue at D=0 (FDR cut off < 0.1%); Normalized enrichment score

(NES). **(C)** Relative expression (RE) of indicated genes by qRT-PCR between matured mini bladder at D=0 (n=3), mini bladders exposed to high (n=3) and low (n=3) solute SHU for D=8. **(D)** Top negatively enriched GO BP pathways ordered by significance in high solute SHU exposed tissue at D=8 vs. control tissue at D=0 (FDR cut off < 0.1%); ). Data represented as mean ± SEM. P-values calculated using one-way Anova with Dunnett's multiple comparison test in **(C)**

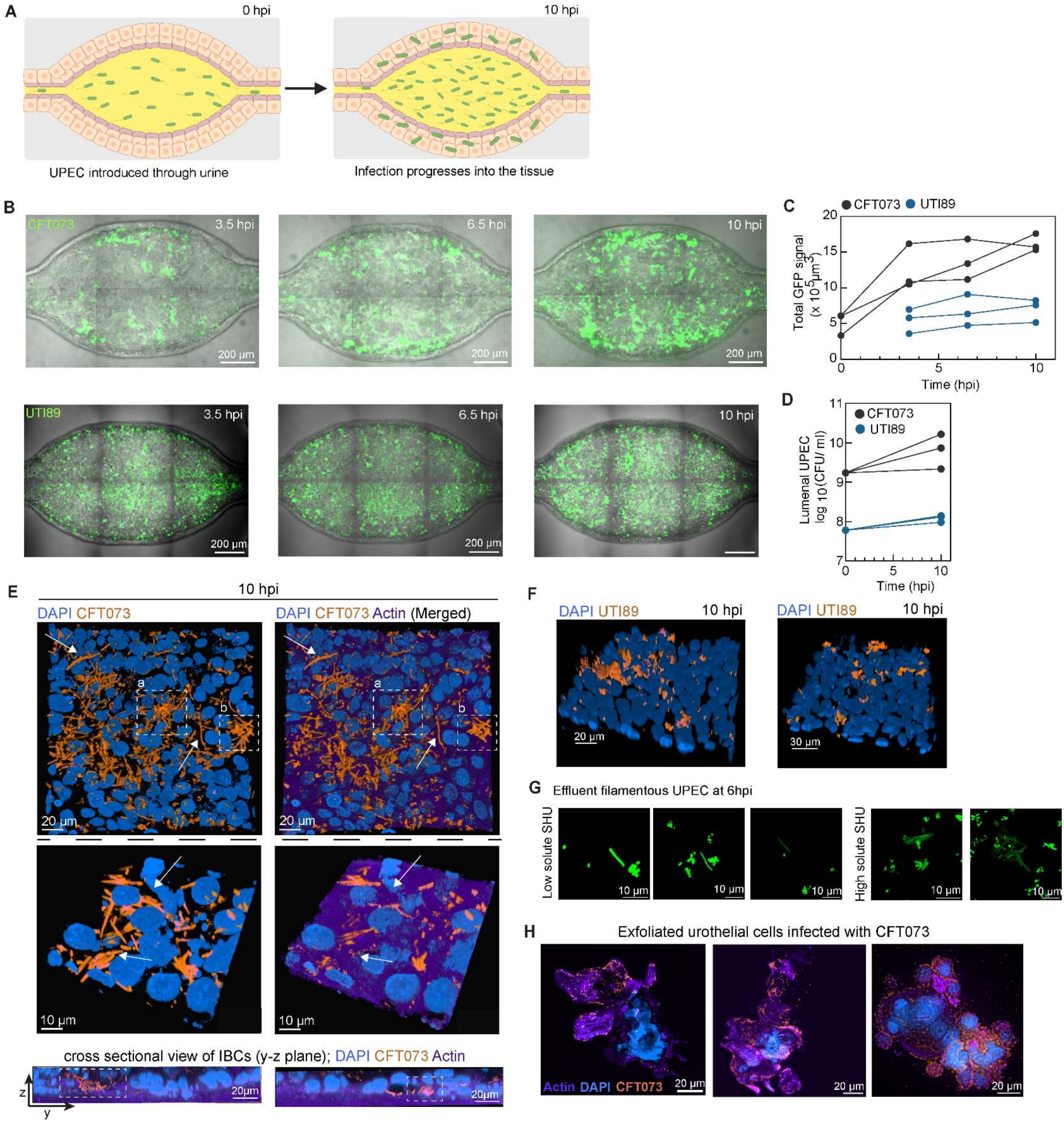


**Figure S6: Gradual progression of infection in mini-bladder tissue during continuous UPEC exposure**

**(A)** Schematic showing infection timeline. **(B)** Snapshots (maximum intensity projection) of timelapse imaging of mini-bladders infected with CFT073 and UTI89 in PHU (UPEC, green). **(C)** Total bacterial volume measured from mini-bladders infected with CFT073 (n=3) and UTI89 (n=3) over time. **(D)**

Urinal CFU collected before and after infection from mini-bladders infected with CFT073 (n=3) and UTI89 (n=3) **(E, F)** Representative 3D view of mini-bladder tissue sections at 10hpi, showing IBCs and filaments of UPEC overlapping with actin stain (actin, purple; DAPI, azure; UPEC, amber; in (E) IBCs marked by white dashed boxes and filaments marked by white arrows). **(G)** Filamentous UPEC in the washout of mini bladders infected with high and low solute SHU (CFT073 UPEC, green) **(H)** Exfoliated urothelial cell clusters infected with CFT073 UPEC collected during washes rel. to Fig 3 (maximum intensity projection) (actin, purple; DAPI, azure; UPEC, amber).

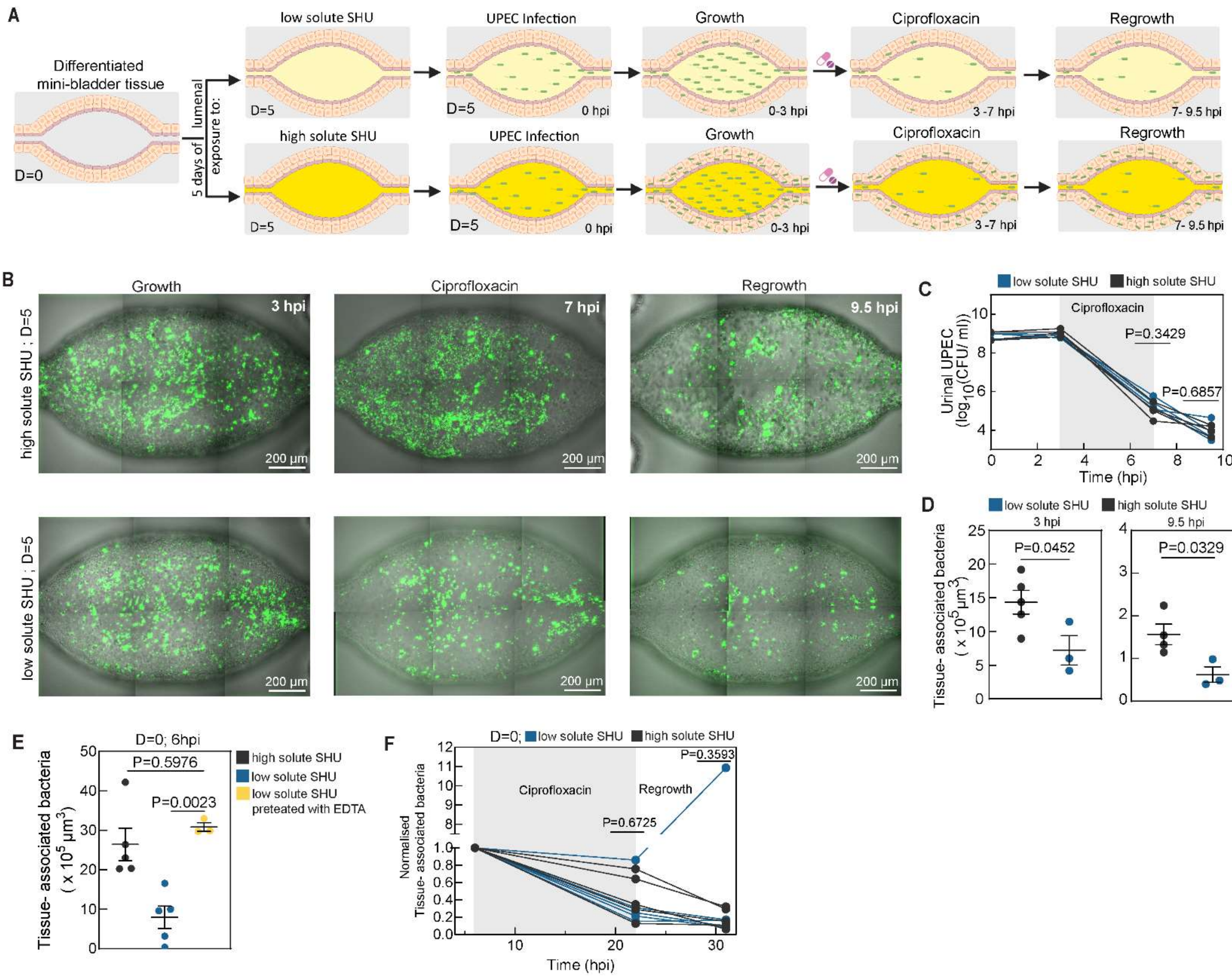


**Figure S7: Long term exposure to high solute SHU lowers antibiotic clearance and increases recurrence**

**(A)** Schematic representation of the timeline of UPEC infection and treatment. **(B)** Snapshots (maximum intensity projection) of timelapse imaging of mini-bladders infected and treated with Ciprofloxacin (100 X MIC in high- solute SHU, 5 μg/mL) in high and low solute SHU at D =5 (UPEC, green). **(C)** Urinal CFU collected through washes from mini-bladders (n=4 each) in high and low solute SHU. **(D)** Tissue associated bacterial volume measured from mini-bladders in high (n=4) and low (n=3) solute SHU **(E)** Tissue associated bacterial volume measured at 6hpi from mini-bladders in high solute (n=5), low solute (n=5), and pre-exposed to EDTA (n=3) on D=0. **(F)** Normalised tissue associated bacterial volume related to Fig 4D. Data represented as mean ± SEM. P-values calculated Mann-Whitney in (**C**), unpaired t test in (**D**), **(F),** one-way Anova with Dunnett's multiple comparison test in (**E**).

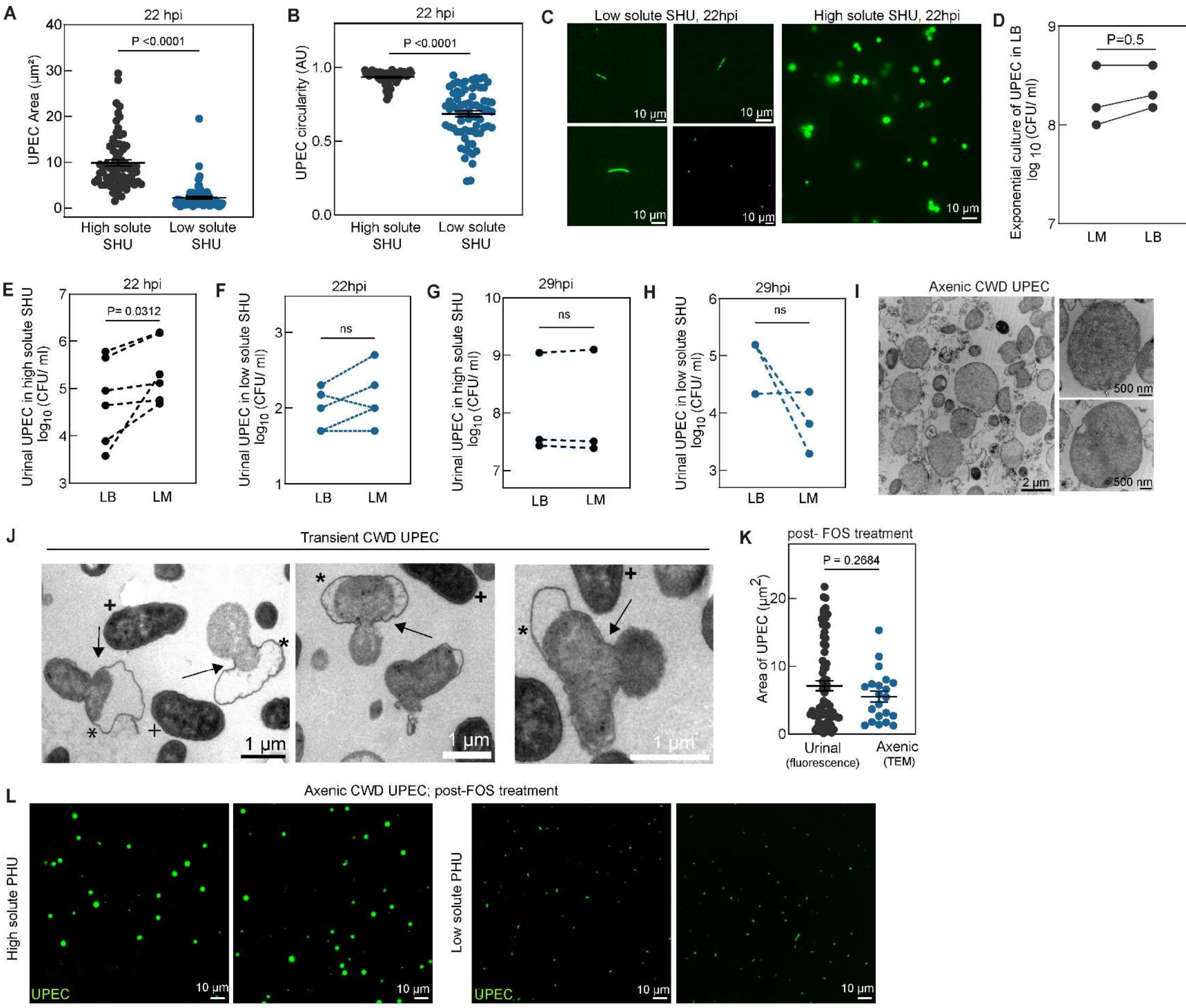


**Figure S8: Characterising urinal UPEC post FOS treatment in high solute SHU**

(**A**) Cross sectional area of urinal UPEC survivors from high (n = 73) and low solute (n= 72) SHU at 22hpi. **(B)** Circularity of urinal UPEC survivors from high (n = 73) and low solute (n= 72) SHU at 22hpi. (**C**) Representative images of urinal UPEC survivors related to panel (A, B) at 22 hpi. **(D)** Comparative survival of exponentially growing axenic UPEC in LB (n=3) **(E, F)** Comparative survival of urinal UPEC samples from high solute (n=6) (E) and low solute (n=5) (F) SHU conditions at 22hpi on LM and LB plates. **(G, H)** Comparative survival of urinal UPEC samples from high solute (n=3) (G) and low solute (n=3) (H) SHU conditions at 29hpi on LM and LB plates. **(I)** TEM images of axenically cultured UPEC treated with Fosfomycin for 16 hours. **(J)** TEM images of UPEC transitioning from rod shaped to CWD form (black plus indicates rod shaped UPEC; black asterisk indicates transitioning UPEC; black arrow highlights the separation of cell wall). **(K)** Cross sectional area of urinal UPEC from Fig 5B (n=72) and axenic UPEC from (**I**) (n=21). **(L)** Representative images of axenic UPEC survivors post Fosfomycin treatment in high and low solute PHU. Data represented as mean ± SEM. P-values calculated using unpaired t test in (**A**), (**B**), (**K**); pairwise Wilcoxon test in **(D),** (**E**), (**F**), (**G**), (**H**).

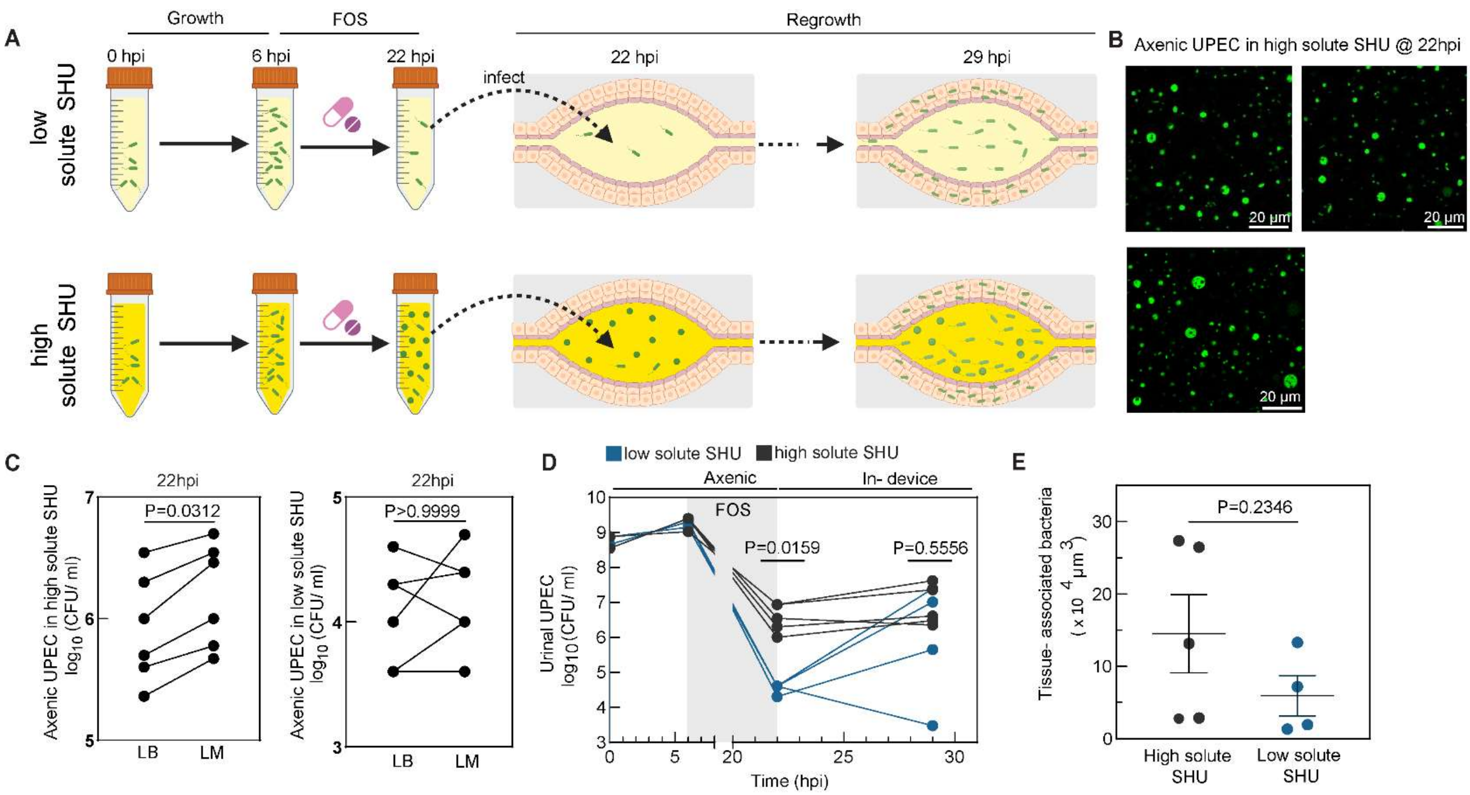


**Figure S9: Presence of tissue associated CWD UPEC is necessary to cause increased recurrence in high solute urine post treatment with Fosfomycin (FOS)**

**(A)** Schematic of infection time course showing initial axenic growth and FOS treatment, followed by infection and regrowth in mini bladder devices under high and low solute SHU. **(B)** Representative images of axenic UPEC in high solute SHU at 22hpi (UPEC, green). (**C**) Comparative survival of axenic UPEC samples from high solute and low solute (n=6 each) urine at 22 hours on LM and LB plates. **(D)** Axenic and urinal CFU collected at different timepoints of infection from mini-bladders in high (n=5) and low (n=4) solute SHU. **(E)** Tissue associated bacterial volume calculated post fixation at 29 hpi, compared between high (n=5) and low (n=4) solute SHU conditions. Data represented as mean ± SEM. P-values calculated using Wilcoxon test in (**C**), Mann-Whitney in (**D**), unpaired t test in (**E**).

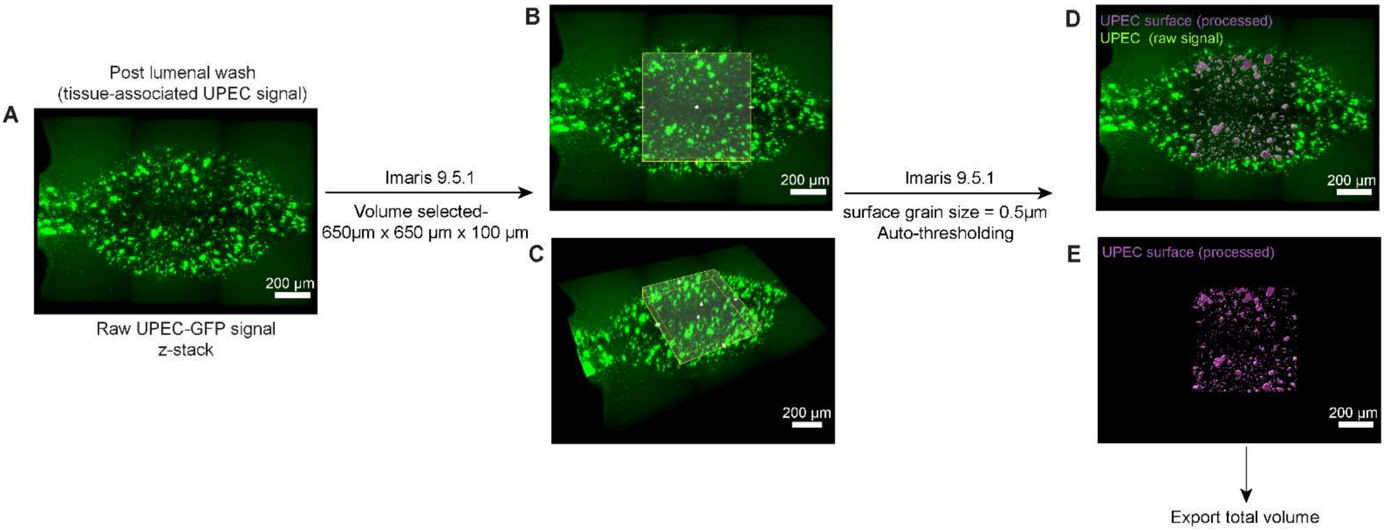


**Figure S10: An illustrated example of the image processing pipeline used to calculate tissue associated bacteria**

**(A-E)** Quantifying tissue associated UPEC from an infected mini-bladder without D-mannose at 3.5 hpi post-wash, representative of Fig 3B. (**A**) The raw z- stack of UPEC-GFP (post-wash) was cropped using a central volume of 650 µm x 650 µm x 100 µm cuboid (**B, C**). The cropped stack was processed using the 'Surface Generation' tool in Imaris with the given parameters (**D, E**). The volume of this generated surface was exported as the value of tissue associated UPEC for the sample.

**Supplementary Table 1**

| | **Osmolarity** (mOsm/kg) | **pH** | **Density** (gm/ml) | **CFT073 Doubling time** (min) | **CFT073 Stationary phase OD** $_{600}$ |
|---|---|---|---|---|---|
| **PHU high concentration** (Sample size = 4) | 663- 664 | 6.8 | 0.99 | 68.7 | 0.292 |
| **PHU low concentration** (Sample size = 4) | 93 - 95 | 7.4 | 0.99 | 117.3 | 0.04 |
| **PHU mid concentration** (Sample size = 3) | 427- 428 | 7.3 | 0.97 | 73.8 | 0.241 |
| **SHU high concentration (temporary buffered aliquots)** | 730 - 740 | 6 | 0.99 | 71.6 | 0.435 |
| **SHU low concentration (temporary buffered aliquots)** | 159 - 192 | 6.4 | 1.01 | 73.4 | 0.437 |

**Supplementary Table 2**

| **Compound** | **Chemical formula** | **Concentration in high solute SHU (mM)** | **Concentration in low solute SHU (mM)** |
|---|---|---|---|
| Sodium chloride | $NaCl$ | 100 | 20 |
| Sodium sulphate | $Na_2SO_4$ | 17 | 3.4 |
| Urea | $CH_4N_2O$ | 280 | 56 |
| Potassium chloride | $KCl$ | 38 | 7.6 |
| Calcium chloride | $CaCl_2$ | 4 | 0.8 |
| Creatinine | $C_4H_7N_3O$ | 9 | 1.8 |
| Citric acid/Trisodium citrate | $Na_3C_6H5O_7$ | 3.4 | 0.68 |
| Ammonium chloride | $NH_4Cl$ | 20 | 4 |
| Magnesium sulphate | $MgSO_4$ | 3.2 | 0.64 |
| Sodium oxalate | $Na_2C_2O_4$ | 0.18 | 0.036 |
| Sodium phosphate monobasic | $NaH_2PO_4$ | 3.6 | 0.72 |
| Sodium phosphate dibasic | $Na_2HPO_4$ | 6.5 | 1.3 |
| Potassium phosphate monobasic | $KH_2PO_4$ | 16 | 3.2 |
| Uric acid | $C5H_4N4O_3$ | 0.6 | 0.12 |
| Sodium bicarbonate | $NaHCO_3$ | 13.5 | 2.7 |
| Magnesium chloride hexahydrate | $MgCl_2$. $6H_2O$ | 3.2 | 0.64 |
| L-(+)-Lactic acid | $C_3H_6O_3$ | 1.1 | 0.22 |
| Iron (II) sulphate heptahydrate | $FeSO_4$. $7H_2O$ | 0.005 | 0.005 |

1 g/L of casamino acids added before adjusting pH to 5.6 using 1M NaOH or 37 % HCl for both high and low solute SHU. This was stored at 4 °C for 1-2 months. Temporary buffered aliquots with additional 20mM HEPES and 1mM $CaCl_2$ were made before all differentiation and infection experiments.

### Supplementary Table 3

| Gene | Primer sequence |
|---|---|
| *UP3A* | Forward:5'- CTCACAGATCCTGAATGCCTACC- 3'<br>Reverse: 5'- CCGTGGACATATTGACCAGGAC- 3' |
| *KRT20* | Forward: 5'-CTG AGG TTC AAC TAA CGG AGC TG-3'<br>Reverse: 5'-AAC AGC GAC TGG AGG TTG GCT A-3' |
| *KRT8* | Forward: 5'-ACA AGG TAG AGC TGG AGT CTC G-3'<br>Reverse: 5'-AGC ACC ACA GAT GTG TCC GAG A-3' |
| *KRT13* | Forward: 5'-GAT GCT GAG GAA TGG TTC CAC G-3'<br>Reverse: 5'-AGC TCC GTG ATC TCT GTC TTG C-3' |
| *P63* | Forward: 5' – CAGGAAGACAGAGTGTGCTGGT- 3'<br>Reverse: 5' – AATTGGACGGCGGTTCATCCCT- 3' |
| *KRT14* | Forward: 5'- TGCCGAGGAATGGTTCTTCACC – 3'<br>Reverse: 5'- GCAGCTCAATCTCCAGGTTCTG- 3' |
| *ZO-1* | Forward: 5'-GTCCAGAATCTCGGAAAAGTGCC – 3'<br>Reverse: 5'- CTTTCAGCGCACCATACCAACC- 3' |

**Movie SMov1**

Time lapse imaging of growth and maturation of a mini-bladder starting from the initial seeding of H-6215 cells. At the end of the movie, a stratified mini-bladder is evident, related to Figure 1.

**Movie SMov2**

Animated Z-stack showing a volume of a differentiated mini-bladder at D = 0 stained with phalloidin (orange) and DAPI (cyan), related to Fig 1B.

**Movie SMov3**

Demonstration of mechanical stretching of a differentiated and stratified mini-bladder.
